# Highly expressed proteins prefer less informative amino acids

**DOI:** 10.1101/054981

**Authors:** Guang-Zhong Wang

## Abstract

The transcriptional and translational systems are essentially information processing systems. However, how to quantify the amount of information decoded during expression remains a mystery. Here, we have proposed a simple method to evaluate the amount of information transcribed and translated during gene expression. We found that although proteins with a high copy number have more information translated, the average number of bits per amino acid is not high. The negative correlation between protein copy number and bits per amino acid indicates the selective pressure to reduce translational errors. Moreover, interacting proteins have similar bits per residue translated. All of these findings highlight the importance of understanding transcription and translation from an information processing perspective.

## Introduction

Recently, there has been a fruitful discussion on the meaning of biological information and how to quantitatively measure the amount of information encoded in the genome (Adami, 2016; Ball, 2016; Barbieri, 2016; Cartwright, 2016; Koonin, 2016; Wills, 2016a, b). As the information encoders (DNA) are evolving, it seems that a simple copy of Shannon’s entropy formula is not plausible at the genomic level, and the meaning of the sequences should be considered (Koonin, 2016). Thus, to quantify information encoded at the genomic level, measurements including the context of sequences need to be developed.

Alternatively, what about measuring the information gain in the transcriptional and translational process? This question is important because it addresses how much information is decoded during expression in certain genomic regions. On the other hand, a major difference between the genetic system and other engineered information systems is that the former is evolving. The information encoded in a specific genomic area changes due to *de novo* mutations and various selective forces. Different parameters (dN, dS, dN/dS ratio, F_op_ and ts/tv ratio) have been developed to describe these processes (Drummond and Wilke, 2008). During the last decade, it has been discovered that expression level is the major factor influencing evolutionary rates, i.e., expression level explains up to 60% of variation in those evolutionary parameters (Drummond et al., 2005; Drummond et al., 2006; Pal et al., 2001). It is believed that widespread anticorrelations exist between coding sequence evolution and expression levels due to selection for reduced translational error, improving translational robustness and avoiding erroneous protein synthesis (Drummond 2008, and Wilke, 2009; Zhang and Yang, 2015). These strong correlations indicate that the genetic information encoded in the genomic DNA is mainly released through decoding activities (transcription and translation).

Instead of directly evaluating the amount of information encoded in the genomic regions, we evaluated the information gain in the process of transcription and translation. We have shown that this parameter varies across different genes and that highly expressed genes prefer to use more informative ribonucleotides and less informative amino acids. Moreover, interacting proteins prefer amino acids with similar bits. All these factors imply the possibility of quantifying information gain and loss in different biological processes.

## Results

### Evaluating the amount of information gained during transcription and translation

We define the amount of information gained per symbol during expression based on how much surprise the decoder machine (RNA polymerase II or ribosomal protein) obtains from this symbol in the intercellular environment. Generally, the lower the concentration of a specific ribonucleotide (A, G, C or U) or amino acid, the greater the surprise when this symbol is transcribed or translated; thus, more information is obtained. For example, the average concentrations (μM) for ribonucleotides in the cell are as follows: ATP: 3,152, GTP: 468, CTP: 278, and UTP: 567 (Traut, 1994). There is a much greater opportunity to capture ATP (0.71) than CTP (0.06) in the cell. The information gained through the transcription of A (-log_2_(0.71) = 0.50 bits) is smaller than through the transcription of C (-log_2_(0.06) = 4.00 bits, see material and methods for more details). In yeast, the translation of cysteine gained the largest amount information (6.2 bits) and lysine the smallest amount (3.7 bits, supplementary table 1). The amount of information (bits) gained for a whole mRNA or protein molecule is the sum of the information of each symbol in the sequence, assuming their transcription and translation are independent. Based on this method, if the compositions of the two genomic regions are similar, the longer the sequence or the higher the copy number of the mRNA or protein molecules, the larger is the amount of information gained during the decoding of this genomic region.

On average, 2.68 bits will be gained per symbol during transcription and ~4.5 bits gained per symbol during translation (Table S1 and Table S2). As the mRNA molecule is more than three times longer than the corresponding protein sequence, more information is obtained through transcription (for yeast, on average 3280 bits gained for mRNA vs. 1899 bits for protein; for mouse, 4921 bits for mRNA vs. 2681 bits for protein). However, if the mRNA and protein copy number are considered, the genetic information translated is significantly higher than the amount transcribed at the genomic scale (Figure 1, p < 2.2E-16 in both comparisons, Wilcoxon signed rank test). As the evolutionary process is the selective usage of genetic information, these observations indicate that more selective pressures exist at the translational level.

**Figure 1.**
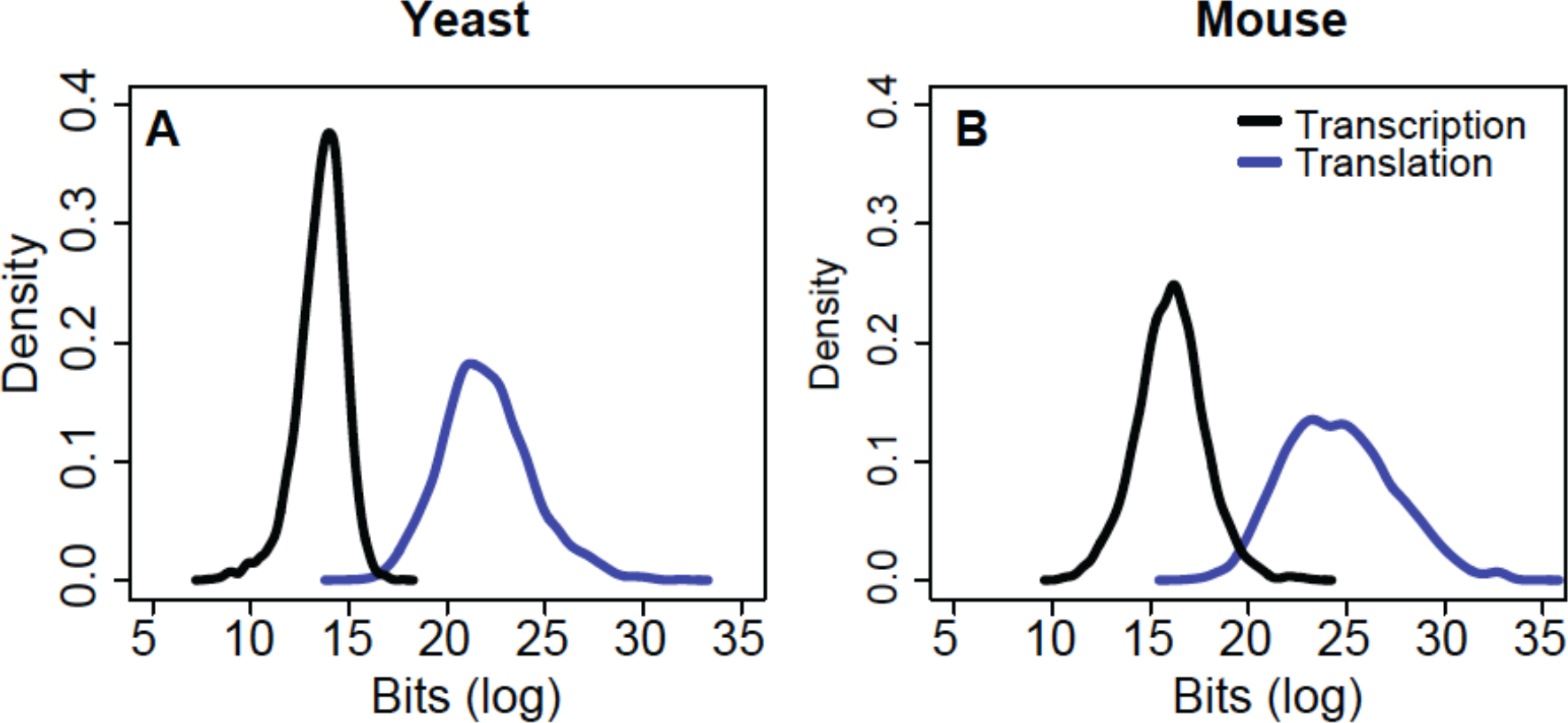
The distribution of total information (bits) transcribed and translated. Although for a single molecule more information is transcribed than translated, the total information gained during translation is significantly higher than during transcription at the genomic scale in both yeast (**A**) and mouse (**B**). Black line, transcription; blue line, translation. Log2 values were used for the x-axis (bits).

### Transcription prefers more informative symbols in highly expressed transcripts

The information gained through both transcriptional and translational activity was estimated at the genomic scale. It has been well established that a gene’s expression level is a major factor influencing genomic evolution, i.e., highly expressed genes evolve slowly (Drummond et al., 2005; Pal et al., 2001). We sought to determine whether there is an association between transcription level and information gain during transcription. We found that the amount of information transcribed is positively correlated with the mRNA copy number (Figures S1A and S1B: for yeast, r = 0.37, p < 2.2E-16 and for mouse, r = 0.72, p < 2.2E-16, Spearman correlation). Similarly, the quantity of average bits gained per symbol is also positively correlated with mRNA copy number (Figure 2A and 2B: for yeast, r = 0.17, p < 2.2E-16 and for mouse, r = 0.12, p < 2.2E-16, Spearman correlation), which means the average uncertainties dismissed are larger in highly expressed transcripts.

**Figure 2.**
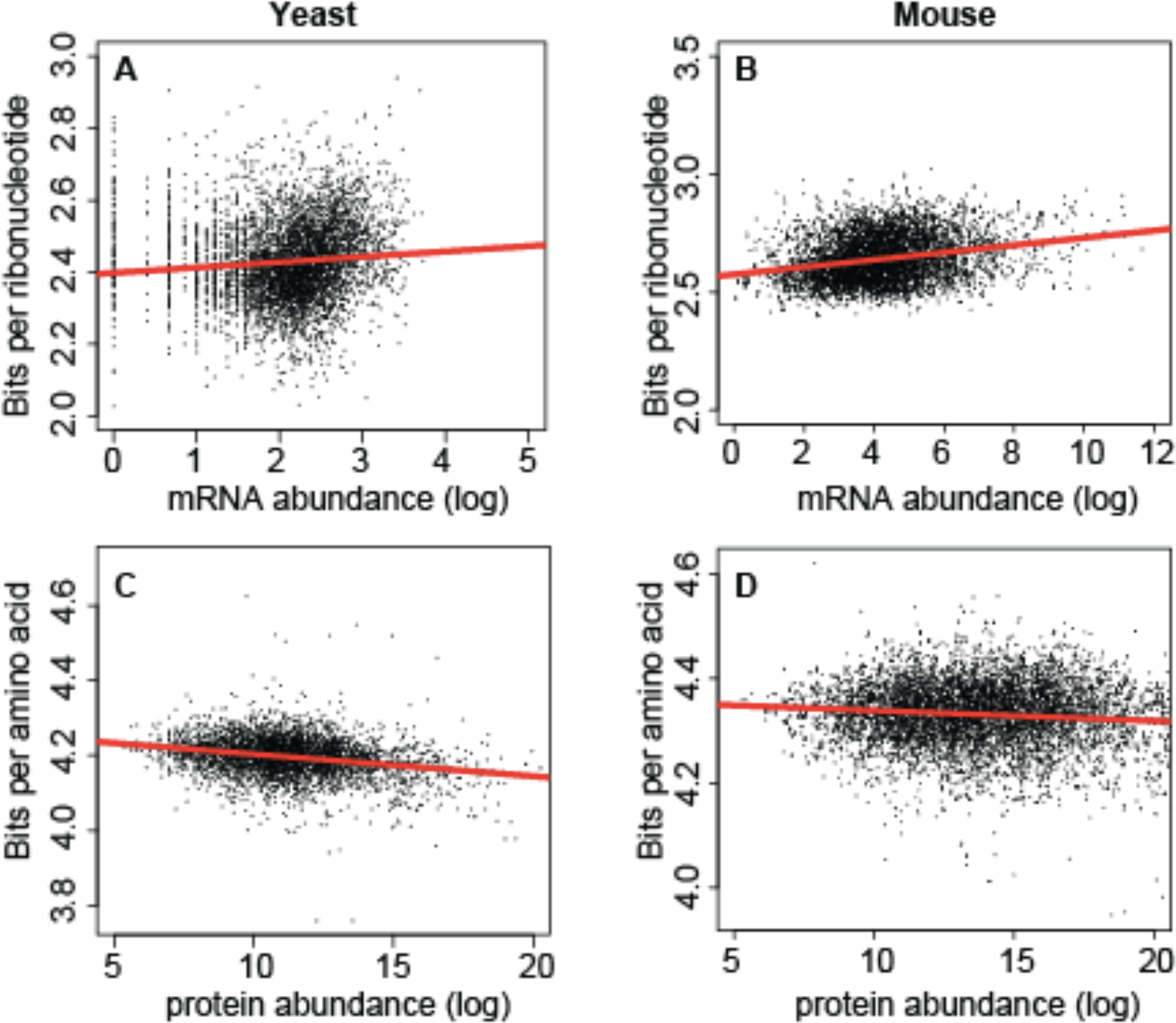
Correlation between mRNA or protein copy number and bits per symbol. Significant positive correlations are observed at transcriptional level (**A** and **B**) and negative correlations are observed at transcription level (**C** and **D**). Red line represents linear regression between abundance and bits per symbol in each plot. Log2 values were used for mRNA and protein abundance.

This observation can be partly explained by the fact that many broadly expressed genes have higher GC content (Lercher et al., 2003), and GC ribonucleotides have higher information content than AU during transcription (CG: 3.25 bits and 4.01 bits *vs*. AU: 0.50 bits and 2.98 bits). Indeed, transcripts with high GC content have significantly higher bits per symbol at the transcriptional level in both yeast and mouse (Figures S2A and S2B: for yeast, r = 0.63, p < 2.2E-16 and for mouse, r = 0.71, p < 2.2E-16, Spearman correlation).

### Transcription prefers more informative symbols in highly expressed transcripts

As similar selective pressures may exist at both the transcriptional and translational levels, we sought to determine whether the positive association between copy number and information gain holds true at the translational level. As shown in figure 2, the information gain during translation is still significantly positively correlated with protein copy number (Figures S1C and S1D: for yeast, r = 0.89, p < 2.2E-16 and for mouse, r = 0.92, p < 2.2E-16, respectively). However, we detected a negative correlation between the information gained per amino acid and protein copy number (Figures 2C and 2D: for yeast, r = -0.23, p < 2.2E-16 and for mouse, r = -0.03, p < 2.2E-16, respectively). The significant correlation remained even after controlling for mRNA copy number and GC content in a linear regression model (p = 2.54E-4 for yeast and p = 1.63E-3 for mouse).

The negative correlation between protein copy number and average bits translated per symbol indicates that highly expressed proteins prefer amino acids with higher concentrations in the intercellular environment, as they are easier to capture (less informative). This finding is consistent with the hypothesis that selection prefers to improve translational accuracy, thus reducing translational errors that will lead to erroneous protein synthesis (Drummond 2009 and Wilke, 2009). Why is this correlation not seen at the transcription level? One possible explanation is that the selection pressure is not strong enough. For example, the probability of capturing the ribonucleotide ATP (71%, the highest among the four ribonucleotides) is almost 10 times higher than the probability of capturing lysine (7.7%, the highest among the twenty amino acids).Moreover, the toxic effect of protein misfolding might be more harmful than the toxic effect of mistakes at the transcriptional level (Drummond 2009and Wilke, 2009).

### Interacting proteins have similar bits translated per symbol

Next, we sought to determine whether interacting genes prefer to utilize building blocks with similar bits. For 10,000 random sampled interacting pairs from the BioGRID database (Stark et al., 2011), the average difference in information gained per amino acid is significantly smaller than expected by chance, both for physically interacting proteins and genetically interacting proteins in yeast (Figures 3A and 3B: mean bits for random pairs: 0.0627+/-0.0006 bits, for physically interacting proteins: 0.05718+/-0.0005 bits, for genetically interacting proteins: 0.0513+/- 0.0004 bits, p <2.2E-16 in both comparisons, Wilcoxon signed rank test). This case also holds in the mouse (Figure 3C: mean bits for random pairs: 0.0682+/-0.0006 bits, for interacting proteins: 0.0583+/-0.0009 bits, p < 2.2E-16, Wilcoxon signed rank test). We validated this observation using 10,000 sampled interacting protein pairs from the STRING database (Szklarczyk et al., 2015) in both yeast and mouse (mean bits for yeast: 0.0565+/- 0.0005 bits, p = 2.0E-10; for mouse: 0.0629+/0.0005 bits, p < 2.2E-16, Wilcoxon signed rank test). Those observations indicate similar selective pressures on the usage of genetic information at the protein level during evolution. At the mRNA level, the difference is still significant for yeast (p < 2.2E-16 in all the comparisons) but not for mouse (p = 0.08 of using BioGRID database and p = 0.35 of using STRING database), which is consistent with the hypothesis that the selective force at the transcriptional level is not as strong as at the translational level.

**Figure 3.**
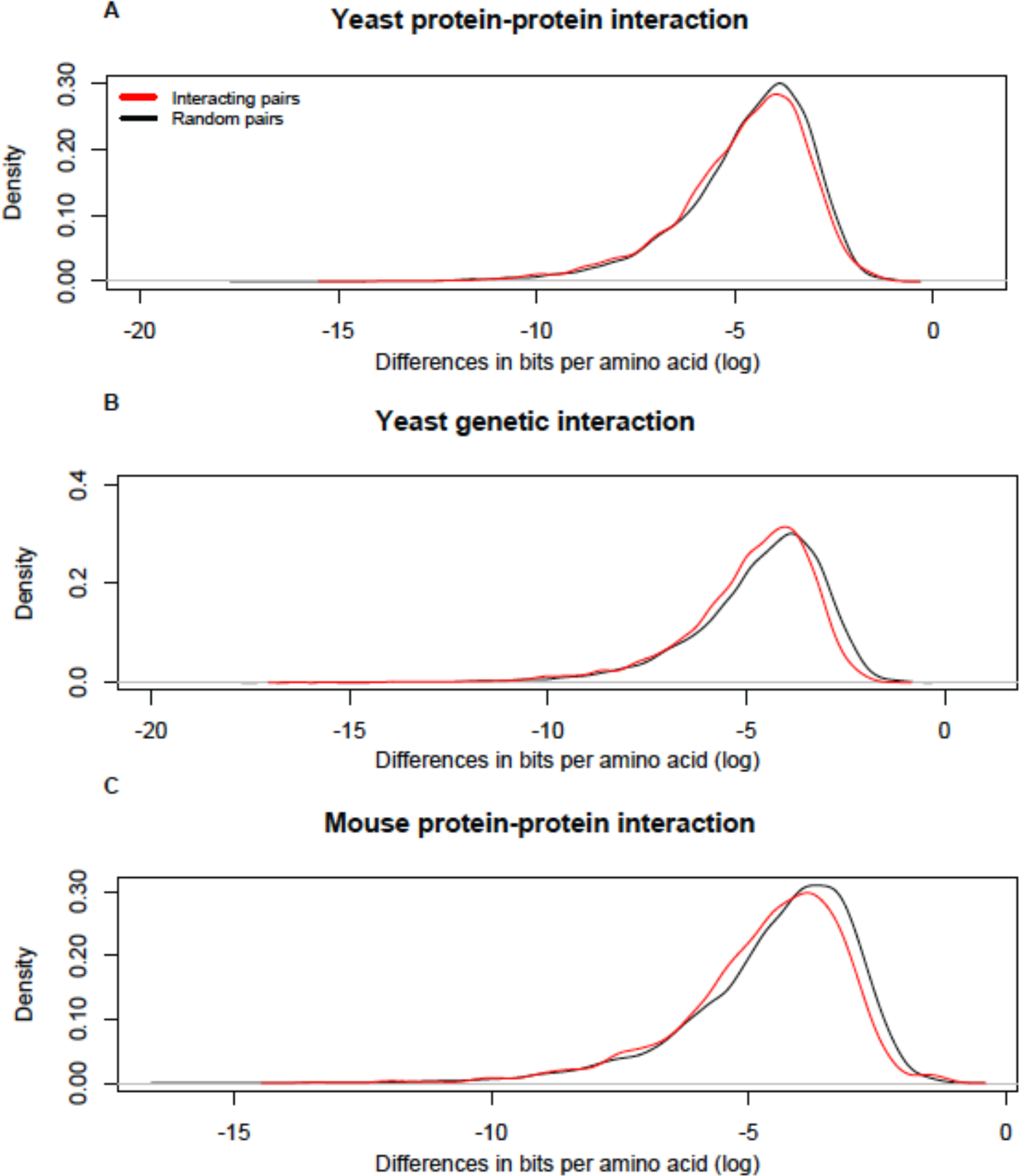
Interacting proteins have similar bits per amino acid translated than expected by chance. The x-axis represents the absolute difference in bits per amino acid between gene pairs (log2). (**A**) yeast protein-protein interaction network; (**B**) yeast genetic interaction network; (**C**) mouse protein-protein interaction network. Red line, interacting gene pairs; black line, random gene pairs.

## Discussion

The decoding of genetic information encoded in the genome depends on gene transcription and translation. By evaluating the amount of uncertainty dismissed per symbol during those processes, we propose a simple method to quantify information gain. This method is based on the probability of obtaining ribonucleotides and amino acids in the intercellular environment. Furthermore, we found that the translational process generates more information than the transcriptional process, both for the average bits per symbol and the total amount of information. Moreover, the average number of bits per symbol is negatively correlated with protein abundance but positively correlated with mRNA abundance. This difference reflects the different selective pressures at those two levels. Interestingly, and consistently with the translational accuracy hypothesis, we have shown that high-abundance proteins prefer to use low-information amino acids, which have higher likelihoods (concentration) of capture in the cell.

In a physically interacting or genetically interacting network, interacting proteins prefer to use amino acids with similar bits, indicating that there are similar selective pressures on the usage of genetic information of such proteins. The differences in bits per symbol between interacting genes and random pairs are not consistent at the translational level, supporting a stronger selective pressure at the translation level. Investigation of the distribution of bits between gene pairs and further in these networks will support the evaluation of the information flow of the interactions.

## Materials and methods

### Genomic sequences

The genomic and protein sequences, together with their annotation files, of yeast and mouse were downloaded from the Ensembl database via the BioMart data mining tool (www.ensembl.org/;version:yeast-EF4,mouse-GRCm38). Short sequences (< 20 amino acids) were excluded for further analysis. For mouse, coding regions, 5’UTR regions, 3’ UTR regions and intron regions were separated the evaluation of information gain. For yeast, we did not distinguish UTR regions and intron regions, as those data are either unavailable or very limited (< 300 introns).

### mRNA and protein copy number

To evaluate the total amount of information gained for each gene, the mRNA and protein copy numbers were considered. The mRNA expression values of yeast were obtained from RNA-seq experiments (Nagalakshmi et al., 2008), and protein copy numbers were obtained from a proteomic study (Ghaemmaghami et al., 2003). For mouse, both mRNA and protein copy number were downloaded from a previous global quantification experiment in the liver (Schwanhausser et al., 2011).

### tRNA copy number

To evaluate the relative probabilities of the available amino acids, we employed the tRNA copy number dataset from the Genomic rRNA Database (http://lowelab.ucsc.edu/GtRNAdb/). All known genomic tRNAs from yeast and mouse were collected, and the tRNAs that do not encode the canonical 20 amino acids were excluded from further analysis.

### Calculating information gain during transcription and translation

We define the amount of information gained per symbol during expression based on the formula *w*(*e_i_*) = −log_2_*p*(*e_i_*), where *p* is the possibility of capturing A, G, C, U or individual amino acids from the intercellular microenvironment (obtained from the concentration of ribonucleotides and the tRNA copy numbers of the 20 amino acids). The number of bits gained from the transcription of a RNA or protein molecule is calculated by 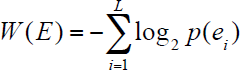, where L is the length of the RNA or protein. Finally, the total amount of information transcribed and translated in a genomic region is evaluated by multiplying the mRNA and protein copy number.

### Interacting protein datasets

Two large-scale datasets were employed for the interacting protein studies: STRING and BioGRID (Stark et al., 2011; Szklarczyk et al., 2015). For yeast, both links from physical interaction and genetic interactions were included, and for mouse, only links from physical interactions were included, as the number of known genetic interactions is too small. For both datasets, 10,000 links were sampled and compared with 10,000 random links obtained from the same gene list. Finally, the absolute differences of bits per symbol were calculated for each gene pair.

### Statistical tests

All statistical tests were performed in R. Correlation analyses were based on Spearman’s methods. The Wilcox rank sum test was used to test the difference in information gain between interacting proteins and random pairs, and the generalized linear regression model was calculated using the “glm” function.

## Contributions

G.–Z. W. conceived the project, performed the analyses and wrote the manuscript.

## Supplementary Information

**Supplementary Figure 1.**
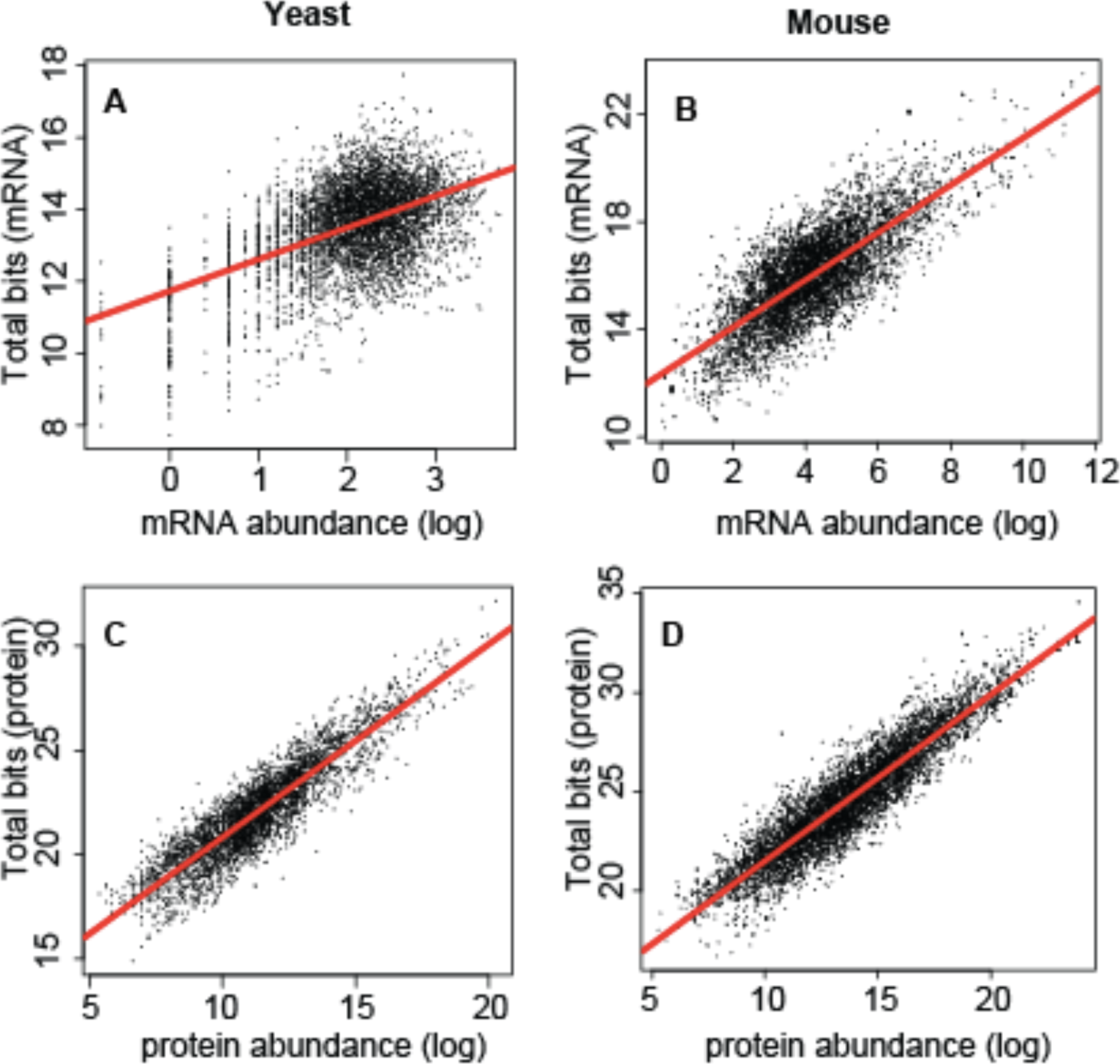
Correlation between mRNA or protein copy number and total bits gained. (**A**) transcription for yeast; (**B**) transcription for mouse; (**C**) translation for yeast; (**D**) translation for mouse. Red line represents linear regression between abundance and bits per symbol in each plot. Log2 values were used for mRNA and protein abundance.

**Supplementary Figure 2.**
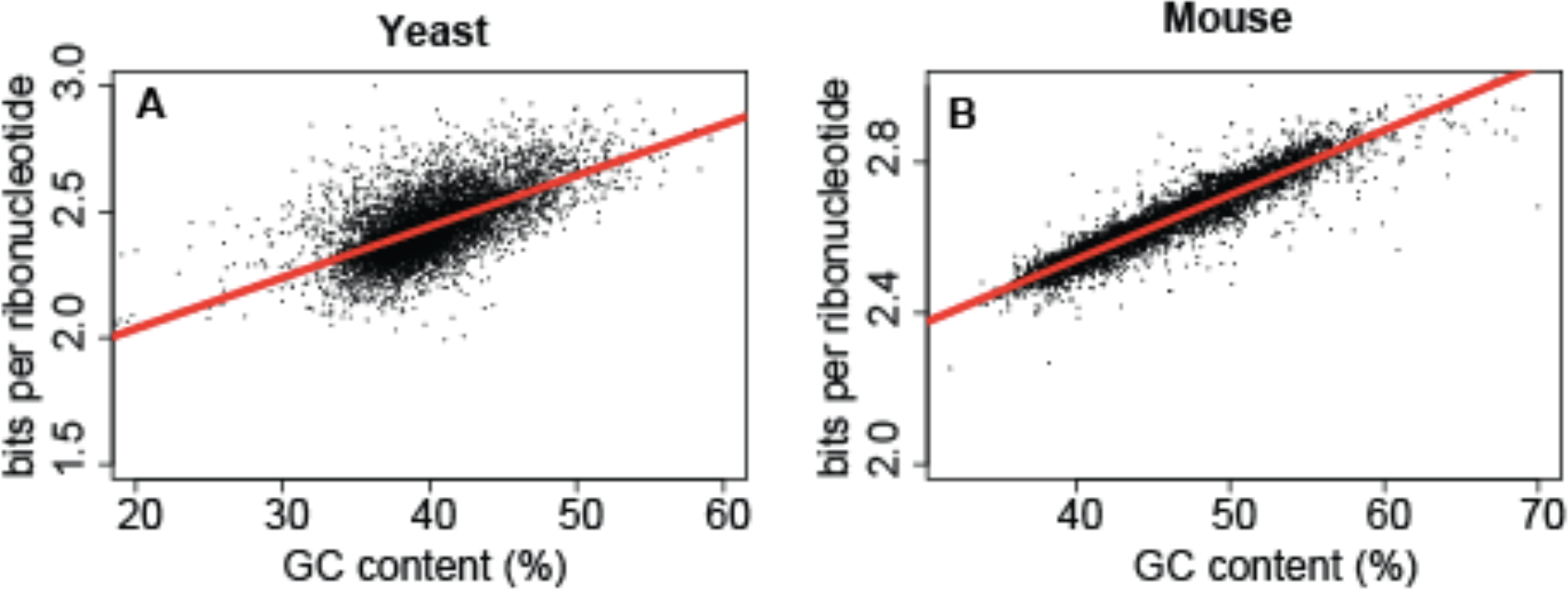
Correlation between GC content (%) and bits gained per symbol. (**A**) yeast; (**B**) mouse. Red line represents linear regression between abundance and bits per symbol.

**Supplementary table 1.** Bits per symbol for the four ribonucleotides.

| Ribonucleotide | Bits |
| --- | --- |
| ATP | 0.50 |
| GTP | 3.25 |
| UTP | 2.98 |
| CTP | 4.01 |

**Supplementary table 2.** Bits per symbol for the twenty amino acids.

| Amino acid | Abbreviation | Bits (yeast) | Bits (mouse) |
| --- | --- | --- | --- |
| Ala | A | 4.07 | 3.43 |
| Gly | G | 3.77 | 3.90 |
| Pro | P | 4.57 | 4.58 |
| Thr | T | 4.07 | 4.58 |
| Val | V | 3.91 | 4.23 |
| Phe | F | 4.70 | 5.95 |
| Asn | N | 4.70 | 4.95 |
| Lys | K | 3.70 | 3.85 |
| Asp | D | 4.16 | 4.75 |
| Glu | E | 4.07 | 4.36 |
| His | H | 5.16 | 5.30 |
| Gln | Q | 4.84 | 4.75 |
| Ser | S | 3.99 | 4.17 |
| Arg | R | 3.77 | 4.17 |
| Leu | L | 3.77 | 3.85 |
| Ile | I | 4.16 | 4.67 |
| Met | M | 4.70 | 4.51 |
| Tyr | Y | 5.16 | 5.43 |
| Cys | C | 6.16 | 2.92 |
| Trp | W | 5.57 | 5.75 |

## References

Adami, C. (2016). What is information? Philos. Trans. A, Math. Phys. Eng. Sci. 374.

Ball, P. (2016). The problems of biological information. Philos. Trans. A, Math. Phys. Eng. Sci. 374.

Barbieri, M. (2016). What is information? Philos. Trans. A, Math. Phys. Eng. Sci. 374.

Cartwright, J.H., Giannerini, S., and Gonzalez, D.L. (2016). DNA as information: at the crossroads between biology, mathematics, physics and chemistry. Philos. Trans. A, Math. Phys. Eng. Sci. 374.

Drummond, D.A., Bloom, J.D., Adami, C., Wilke, C.O., and Arnold, F.H. (2005). Why highly expressed proteins evolve slowly. Proc. Natl. Acad. Sci. USA 102, 14338–14343.

Drummond, D.A., Raval, A., and Wilke, C.O. (2006). A single determinant dominates the rate of yeast protein evolution. Mol. Biol. Evol. 23, 327–337.

Drummond, D.A., and Wilke, C.O. (2008). Mistranslation-induced protein misfolding as a dominant constraint on coding-sequence evolution. Cell 134, 341–352.

Drummond, D.A., and Wilke, C.O. (2009). The evolutionary consequences of erroneous protein synthesis. Nat. Rev. Genet. 10, 715–724.

Ghaemmaghami, S., Huh, W.K., Bower, K., Howson, R.W., Belle, A., Dephoure, N., O'Shea, E.K., and Weissman, J.S. (2003). Global analysis of protein expression in yeast. Nature 425, 737–741.

Koonin, E.V. (2016). The meaning of biological information. Philos. Trans. A, Math. Phys. Eng. Sci. 374.

Lercher, M.J., Urrutia, A.O., Pavlicek, A., and Hurst, L.D. (2003). A unification of mosaic structures in the human genome. Hum. Mol. Genet. 12, 2411–2415.

Nagalakshmi, U., Wang, Z., Waern, K., Shou, C., Raha, D., Gerstein, M., and Snyder, M. (2008). The transcriptional landscape of the yeast genome defined by RNA sequencing. Science 320, 1344–1349.

Pal, C., Papp, B., and Hurst, L.D. (2001). Highly expressed genes in yeast evolve slowly. Genetics 158, 927–931.

Schwanhausser, B., Busse, D., Li, N., Dittmar, G., Schuchhardt, J., Wolf, J., Chen, W., and Selbach, M. (2011). Global quantification of mammalian gene expression control. Nature 473, 337–342.

Stark, C., Breitkreutz, B.J., Chatr-Aryamontri, A., Boucher, L., Oughtred, R., Livstone, M.S., Nixon, J., Van Auken, K., Wang, X., Shi, X., et al. (2011). The BioGRID Interaction Database: 2011 update. Nucleic Acids Res. 39, D698–704.

Szklarczyk, D., Franceschini, A., Wyder, S., Forslund, K., Heller, D., Huerta-Cepas, J., Simonovic, M., Roth, A, Santos, A., Tsafou, K.P., et al. (2015). STRING v10: protein-protein interaction networks, integrated over the tree of life. Nucleic Acids Res. 43, D447–452.

Traut, T.W. (1994). Physiological concentrations of purines and pyrimidines. Mol. Cell Biochem. 140, 1–22.

Wills, P.R. (2016a). DNA as information. Philos. Trans. A, Math. Phys. Eng. Sci. 374.

Wills, P.R. (2016b). The generation of meaningful information in molecular systems. Philos. Trans. A, Math. Phys. Eng. Sci. 374.

Zhang, J., and Yang, J.R. (2015). Determinants of the rate of protein sequence evolution. Nat. Rev. Genet. 16, 409–420.

